# A 22,403 marker composite genetic linkage map for cassava (*Manihot esculenta* Crantz) derived from ten populations

**DOI:** 10.1101/010637

**Authors:** International Cassava Genetic Map Consortium (ICGMC).

## Abstract

Cassava (*Manihot esculenta* Crantz) is a major staple crop in Africa, Asia, and South America, and its starchy roots provide nourishment for 800 million people worldwide. Although native to South America, cassava was brought to Africa approximately 400 years ago and is now widely cultivated across sub-Saharan Africa. The widespread use of clonal planting material, however, aids the spread of disease. Breeding for disease resistance and improved yield began in the 1920s and has accelerated in the last 45 years. To assist in the rapid identification of markers for pathogen resistance and crop traits, and to accelerate breeding programs, we generated a framework map for *M. esculenta* Crantz derived from reduced representation sequencing (genotyping-by-sequencing [GBS]). The composite 2,412 cM map integrates ten biparental maps (comprising 3,480 meioses) and organizes 22,403 genetic markers on 18 chromosomes, in agreement with the observed karyotype. The map anchors 71.9% of the draft genome assembly and 90.7% of the predicted protein-coding genes. The resulting chromosome-anchored genome sequence provides an essential framework for identification of trait markers and causal genes as well as genomics-enhanced breeding of this important crop.

## INTRODUCTION

Cassava (*Manihot esculenta* Crantz) is cultivated as a staple in much of Africa, South America and Asia as it is easy to grow with limited inputs (HOWELER *et al.* 2013). Smallholder farmers typically grow cassava in plots of a hectare or less. Its starchy roots can be left in the ground until they are needed, making cassava an excellent food security crop. Since cassava can be clonally propagated, desirable varieties can be genetically fixed immediately and multiplied for distribution. The crop is relatively drought tolerant, and hence likely robust to climate change. Cassava is also grown for industrial starch production and biofuel applications, particularly in Southeast Asia (HOWELER *et al.* 2013).

Despite its advantages, cassava faces several biotic and abiotic challenges. Clonal propagation facilitates the rapid spread of bacterial and viral diseases. Furthermore, roots of most farmer-preferred varieties are nutrient poor and deteriorate rapidly post-harvest, preventing farmers from generating income from the sale of excess crop.

The use of modern genetic and genomic techniques, such as quantitative trait locus (QTL) mapping, genomic selection, genome-wide association studies (GWAS), and genetic engineering, can accelerate the pace of disease resistance locus identification and trait improvement. However, these require a high-quality genome assembly and a dense genetic map. A draft cassava genome assembly was generated and covers 532.5 Mb (69%) of the estimated 770 Mb cassava genome (AWOLEYE *et al.* 1994). This assembly captures half of the genome sequence in the 487 largest scaffolds, all longer than 258 kb, and 90% of the assembly is accounted for in 2,654 scaffolds all longer than 23 kb; however these are not linked to chromosomes (PROCHNIK *et al.* 2012). To date, a number of genetic maps have been generated for cassava using different marker systems: restriction fragment length polymorphism (RFLP), random amplified polymorphic DNA (RAPD), microsatellite, and isoenzyme (FREGENE *et al.* 1997); simple sequence repeat (SSR) (OKOGBENIN *et al.* 2006); amplified fragment length polymorphism (AFLP) and SSR (KUNKEAW *et al.* 2010); expressed sequence tag (EST) and EST-SSR (SRAPHET *et al.* 2011); SSR and EST-derived single nucleotide polymorphism (SNP) (RABBI *et al.* 2012); and GBS-SNP (RABBI *et al.* 2014a; RABBI *et al.* 2014b). These maps have low marker resolution and/or do not resolve a complete set of linkage groups (LGs) representative of the 2n = 36 karyotype of cassava (DE CARVALHO AND GUERRA 2002). Furthermore, the densest GBS-derived SNP map anchors only 313.3 Mb (58.7%) of the reference genome assembly (RABBI *et al.* 2014b). Since there are many cassava breeding programs, working with varying accessions and trait(s) of interest, a broadly useful genetic map should include markers that segregate in diverse populations.

A single biparental cassava cross rarely yields enough progeny to make a dense map, and in any event, would only capture markers from a small sample of haplotypes segregating in the species. We therefore merged ten maps derived from diverse parents to produce a composite genetic map. In order to generate such a composite map, we obtained one S1 and nine F1 populations (fourteen parents total) from African cassava breeding projects. Markers were generated via GBS (ELSHIRE *et al.* 2011) and a map was constructed from each of the ten crosses with JoinMap (VAN OOIJEN 2011). These maps were merged with LPmerge (ENDELMAN AND PLOMION 2014) to generate a 2,412 centiMorgan (cM) genetic map comprising 18 LGs, in agreement with the number of chromosomes found cytogenetically (DE CARVALHO AND GUERRA 2002). Furthermore, 71.9% of the genome assembly was anchored to the genetic map. The resulting chromosome-scale assembly will accelerate the application of a wide variety of modern tools for crop improvement.

## MATERIALS AND METHODS

### Generation of mapping populations

Nine biparental (F1) and one self-pollinated (S1) crosses were performed (Table 1). Biparental populations NxA, KAR, NCAR, MT, NDLAR and ARAL were generated from crossing blocks planted in four locations in Tanzania: Naliendele (10°23' S, 40°09' E, altitude ∼137 m above sea level [ASL]), Kibaha (6°46' S, 38°58' E, ∼162 m ASL), Ukiriguru (2°43'23” S, 33°1'39” E, 1229 m ASL) and Maruku (1°24'55” S; 31°46'48” E, 1340 m ASL). Populations 412x425, MP4 and MP5 were generated at International Institute of Tropical Agriculture (IITA) crossing sites in Nigeria. Pedigrees of parents are shown in Table 2. Cassava stakes for planting were obtained from research stations or farmers’ fields. For populations developed in Tanzania (with the exception of ARAL) more than 4,000 hand pollinations were performed according to Kawano (KAWANO 1980). Over 10,000 seeds were generated with over 1,000 seeds per population except for population MT. After three months, seeds were sown in trays and raised in a screen house for one month before transplanting into the field at Makutupora research station in Dodoma, Tanzania. Poor germination rates resulted in a total of approximately 3,500 seedlings, with further losses incurred during field establishment.

**Table 1.** Mapping populations used in this study. Nine biparental (F1) and one self-pollinated (S1) populations were generated in which a variety of disease and agronomic traits were segregating. After sequencing, individuals that were not full sibs and/or had insufficient read depth for accurate variant calling were removed prior to map construction.

| Population | Female parent | Male parent | Cross type | Number of individuals sequenced | Number of validated progeny | Purpose of cross (segregating traits) |
| --- | --- | --- | --- | --- | --- | --- |
| ARAL<sup>a</sup> | AR40-6 | Albert | F1 | 154 | 129 | Cassava brown streak disease (CBSD) and green mite resistance |
| KAR<sup>a</sup> | Kiroba | AR37-80 | F1 | 192 | 132 | CBSD and green mite resistance |
| MP4<sup>b</sup> | TMS-IBA30001 | TMS-IBA961089A | F1 | 190 | 177 | Starch, dry matter content, cassava mosaic disease (CMD) resistance, and root rot |
| MP5<sup>b</sup> | TMS-IBA961089A | TMS-IBA30001 | F1 | 187 | 162 | Starch, dry matter content, CMD resistance, and root rot |
| MT<sup>a</sup> | Mkombozi | Unknown | F1 | 157 | 135 | CBSD resistance |
| NCAR<sup>a</sup> | Nachinyaya | AR37-80 | F1 | 240 | 233 | CBSD and green mite resistance |
| NDLAR<sup>a</sup> | NDL06/132 | AR37-80 | F1 | 247 | 244 | CBSD and green mite resistance |
| NxA<sup>a</sup> | Namikonga | Albert | F1 | 303 | 256 | CBSD resistance |
| TMEB419-S1<sup>c</sup> | TMEB419 | TMEB419 | S1 | 149 | 117 | Starch content |
| 412x425<sup>b</sup> | TMS-IBA4(2)1425 | TMS-IBA011412 | F1 | 177 | 155 | Root carotenoid content, CMD resistance |

<sup>a</sup> Ferguson lab, IITA Nairobi, Kenya
<sup>b</sup> Rabbi lab, IITA Ibadan, Nigeria
<sup>c</sup> Egesi lab, National Root Crops Research Institute (NRCRI) Umudike, Nigeria

**Table 2.** Provenance and pedigrees of parents of mapping populations.

| Accession name | Provenance and/or pedigree information |
| --- | --- |
| Namikonga | Known as 'Kaleso' in Kenya. Third backcross from inter-specific hybrid (46106/27) from *M. glaziovii* from Amani breeding program (JENNINGS 1960; HILLOCKS AND JENNINGS 2003). |
| AR40-6 | A CIAT cross between a CMD resistant variety C39 from IITA and CW259-42. CW259-42 is a backcross of MTAI 8 (Rayong 60) and an interspecific cross between *M.e. ssp flabellifolia* and CM 2766-5. |
| Kiroba | Farmer variety from Tanzania. |
| NDL06/132 | Breeding line selected at Agricultural Research Institute (ARI) Naliendele in southern Tanzania. It is an S1 of variety NAL90/34 which showed strong resistance to CBSD (HILLOCKS AND JENNINGS 2003) and is half sib of Kibaha, which has *M.e. ssp. flabellifolia* background. |
| Mkombozi | It is a half sib of 92/0099S2(SM). 92/0099 is a cross 91934 x 81/00032. 91934 is 58308 x Ogunjobi, and 81/00032 is a cross between U/1421 and P-2. Also known as MM96/4684. 58308 is from Amani material selected at Moore Estate, and thus has wild species introgression in its pedigree. |
| Nachinyaya | Farmer variety from Tanzania. |
| Albert | Farmer variety from Tanzania. |
| TMS-IBA4(2)1425 | 58308 x Oyarugba Funfun. |
| AR37-80 | A CIAT cross between a CMD resistant line (C33) from IITA and CW259-42. |
| TMEB419 | TMEB419 is a clone collected in Togo with an unknown pedigree, and released as a variety in Nigeria in 2005 with the name TME419. It is also known by the name Gbasekoute. TMEB419 is very popular all over Nigeria and many other African countries where it has been introduced because it has high dry matter and starch content. The quality of food products from it such as gari and fufu are excellent. It also has erect stems with minimal branding, which facilitates intercropping as well as higher planting densities. |
| TMS-IBA011412 | Cloned in 2001, this is an improved IITA variety with resistance to CMD and it accumulates provitamin A carotenoids in its storage roots. It is the progeny of a cross between TMS-IBA950971 and TMS-IBA940561. |
| TMS-IBA30001 | One of the early TMS series of clones derived from early CMD resistance breeding work in West Africa. It traces its ancestry to variety 58308 which came from interspecific crosses with *M. glaziovii*. |
| TMS-IBA961089A | TMS-IBA961089A is an improved variety from IITA that shows strong resistance to CMD that is likely to be a result of its parentage. Its female parent is TMS-MOK940461, a half-sib of Nigerian landrace TMEB9, which possesses qualitative resistance to CMD. Its other parent, TMS-IBA9001554, is a half-sib of TMS-IBA30572, an improved variety cloned in 1973, with quantitative resistance to CMD that is derived from variety 58308, a hybrid derived directly from recombination of the *M. glaziovii* x *M. esculenta* triple-backcrosses. |

For the TMEB419 S1 cross, cassava stakes for planting were derived from the NRCRI cassava breeding experimental plots and planted at a hybridization plot at Ubiaja, Nigeria. Up to 5,000 hand pollinations were performed by selfing the same variety using it as both male and female parents but with flowers and pollen from different plants. About 800 seeds were generated. Seeds were sown in trays and raised in a screen house for one month before transplanting into the field at NRCRI Umudike experimental field in Nigeria. Poor germination rates resulted in a total of approximately 200 seedlings, with further losses incurred during field establishment.

### DNA isolation

Genomic DNA was isolated from the F1 populations according to Dellaporta *et al.* (1983), with some modifications to allow for processing of many samples in parallel in small volumes. DNA was extracted from population TMEB419-S1 using DNeasy Plant Mini Kit (Qiagen) following manufacturer protocol. In all cases, young apical leaves were first freeze-dried then ground using a GenoGrinder beadmill at 1,500 strokes/min for 2 min. NxA, KAR, NCAR, MT, NDLAR and ARAL populations were initially screened as in Kawuki *et al.* (2013) with approximately 12 SSR markers that were polymorphic among the parents, to detect off-types and selfs. These were removed from the population prior to GBS. Additional off-types, half-sibs, and selfs were later detected by GBS and removed from mapping populations (see below).

### Genotyping-by-sequencing library preparation and sequencing

GBS library construction was performed at the University of California, Berkeley (UC Berkeley) using a protocol adapted from Elshire *et al.* (2011). The restriction enzyme *Ape*KI [New England Biolabs (NEB)] was used in conjunction with the barcode sequences included in the Elshire paper. Differences from their protocol following protocol optimization are described below and discussed further in the Results and Discussion section. Y-shaped adapters were designed based on the Y-shaped Illumina DNA paired-end (PE) adapters (Illumina, Inc.; Figure S1). “Forward” adapter oligos had the sequence 5’ ACACTCTTTCCCTACACGACGCTCTTCCGATCTxxxx, and “reverse” oligos the sequence 5’ CWGyyyyAGATCGGAAGAGCGGTTCAGCAGGAATGCCGAG; where xxxx represents the 4–8 bp barcode and yyyy its reverse complement. CWG is the *Ape*KI-specific overhang. Reverse oligos were phosphorylated at the 5’ end. Adapters were ordered from IDT as a “primer premix plate”, with standard desalting, in a total volume of 50 μl with each oligo at a concentration of 200 μM. Adapters were annealed at 50 μM in TE using a thermocycler: 95°C 4 min, ramp –70°C at 0.1°C/5 sec, 25°C 5 min, hold at 4°C. Annealing was confirmed by running adapters on a 4% agarose gel next to single-stranded oligos of similar length, as annealed adapters run at a larger size. Annealed adapters were diluted 1:10 and then to 12 ng/μl in TE, and finally to 1.25 ng/μl in 10 mM Tris pH 8. Adapter volumes for the last two dilutions were based on quantitation with Picogreen reagent (Invitrogen) and an FLx800 microplate reader (Biotek Instruments, Inc.).

Libraries comprised 63–96 samples. Sample DNA quantitation was performed with Picogreen. Typically, DNA samples were diluted in water to approximately 20 ng/μl and re-quantitated before library preparation. Digests were performed on a 96-well plate, and each consisted of 100 ng DNA in 20 μl 1× NEBuffer #3 (NEB) with 5 U *Ape*KI. Three microliters of the desired pre-annealed and diluted adapter was added to each well of digested DNA, followed by ligation mix containing 720 cohesive end units T4 DNA ligase and 5 μl T4 DNA Ligase Reaction Buffer (NEB), to a final ligation volume of 50 μl. Ligations were pooled such that each offspring sample contributed an equal amount of DNA. For parental DNA samples, in order to ensure adequate sequence coverage a greater amount of digested/ligated DNA was added to the pool. Pools contained a total amount of 1–2 μg DNA, and were purified and concentrated using the MinElute PCR Purification Kit (Qiagen).

Size selection was performed on pooled libraries using a 2% agarose gel run at 140 V. A size fraction of 400–800 bp was excised from the gel and purified via the MinElute Gel Extraction Kit (Qiagen), melting the gel at room temperature to avoid G-C bias (QUAIL *et al.* 2008). Most libraries were amplified with five PCR cycles using Phusion polymerase (NEB), 460 ng of each PCR primer, and an extension time of 45 seconds. After PCR amplification, libraries were cleaned using 0.7 volumes AMPure XP SPRI beads (Beckman Coulter, Inc.). Size distributions of Illumina libraries were assayed by the Vincent Coates Genomic Sequencing Laboratory (VCGSL) at UC Berkeley using a 2100 Bioanalyzer (Agilent Technologies, Inc.). Library concentrations were determined by the VCGSL using a Qubit (Life Technologies) and quantitative PCR. Quantitative PCR was performed with Kapa Biosystems’ Illumina Library Quantification Kits and and Roche LightCycler 480, following all kit protocols. One hundred basepair paired-end sequencing was performed by the VCGSL on HiSeq 2000 or 2500 instruments (Illumina, Inc.). Some libraries were sequenced more than once, usually because the first run was suboptimal. Sequence read totals for each population are given in Table 3.

**Table 3.** Sequence reads and variability. Summary statistics are shown for the ten mapping populations used in this study. The coefficient of variation for a given population is an average of libraries sequenced for that population. The total (for reads) or average (for average reads per barcode and coefficient of variation) are shown in the last line (bold).

| Population name | Total reads | Average number of reads / barcode | Coefficient of variation |
| --- | --- | --- | --- |
| ARAL | 1,004,686,146 | 6,440,295 | 0.3689 |
| KAR | 913,517,846 | 3,713,487 | 0.3669 |
| MP4 | 734,460,036 | 3,672,300 | 0.6334 |
| MP5 | 735,540,482 | 3,733,708 | 0.7023 |
| MT | 609,010,716 | 3,421,408 | 0.7470 |
| NCAR | 1,076,417,780 | 4,375,682 | 0.3830 |
| NDLAR | 1,329,433,056 | 5,113,204 | 0.3741 |
| NxA | 1,189,124,090 | 3,823,550 | 0.4403 |
| TMEB419-S1 | 693,520,182 | 4,532,811 | 0.3051 |
| 412x425 | 1,183,937,274 | 6,399,660 | 0.5171 |
| **Total or average** | **9,406,878,938** | **3,770,292** | **0.4853** |

### Genotype calling and filtering

GBS data were analyzed by pipelining several widely used sequence analysis tools with custom scripts to extract markers with parental genotype combinations useful for the cross-pollinated (CP) genetic mapping strategy implemented by JoinMap (v4.1 2013, July 11 release) (VAN OOIJEN 2011). An outline of the pipeline is shown in Figure 1 and step-by-step command-line instructions are available at https://bitbucket.org/rokhsar-lab/gbs-analysis.

**Figure 1.**
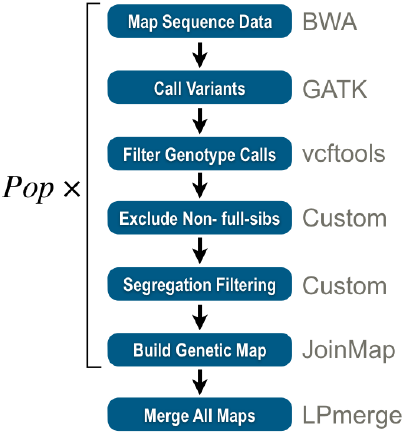
Data analysis pipeline used in this study. Using a combination of publicly available and custom tools (gray text), the pipeline starts with sequence data and generates a map for to each population (Pop) through a series of analyses (white text on blue). Lastly, the maps are merged using LPmerge to generate a single composite map (ENDELMAN AND PLOMION 2014).

To ensure high-quality data for genetic map estimation, the raw read data were trimmed of adapter sequences with the fastq-mcf tool from the ea-utils package (ARONESTY 2011), demultiplexed with an allowance of one mismatch in the barcode sequence using a custom script, and then base-quality trimmed (Q=28) and aligned to the cassava v4.1 draft genome assembly using BWA (LI AND DURBIN 2009). Variants and genotypes were called using the HaplotypeCaller tool from the GATK (v2.7-2) (MCKENNA *et al.* 2010) with a minimum mapping quality threshold of 25. As map-estimation software can be sensitive to excessive missing data and genotyping error, genotypes with quality scores and read depths less than 30 and 10x, respectively, were marked as missing data and sites with over 20% missing genotypes were discarded. To avoid spurious genotype calls within repeat regions, sites with average depth greater than approximately 120 reads per individual or with log_10_(GATK HaplotypeScore+1) values greater than 0.5 were removed. A maximum log_10_(GATK HaplotypeScore+1) value of 1.0 was enforced for the ARAL and 412x425 populations because they had been sequenced more deeply.

Further filtering removed individuals with either insufficient data or half-sib, off-type, or (for biparental crosses) self-pollinated individuals (see below). The chi-squared P-value for F1 Mendelian ratios was then calculated for each variant site, and sites with P < 0.05 were discarded. Parental genotypes were then inferred from the segregation pattern of each marker in the progeny and this information was used to impute missing parental genotypes in either parent. Markers were then grouped into non-overlapping 50 kb bins and one marker of each segregating type (*i.e.*, lm x ll, nn x np, hk x hk, ef x eg, and ab x cd) with the most genotyped individuals was chosen to represent each bin.

### Identification of off-types, half-sibs, and selfs in the progeny

Cassava farmers prefer to grow several varieties in a field at one time and this often leads to volunteer seedlings and hence genetic variability within nominally clonally propagated cassava. This can create spurious genetic variation within varieties used to generate populations. We therefore checked the progeny from each biparental mapping population for the presence of off-types, half-sibs and selfs. We used metrics of relatedness to determine whether a progeny was a full-sib F1 (or S1). First, we used the frequency of genotypes violating, or inconsistent with, the expected Mendelian patterns of segregation. We defined this as the fraction of genotypes homozygous for the minor/unshared parental allele (or contained non-parental alleles) per individual. Individuals with a rate of Mendelian inconsistency in excess of 0.005 at a minimum genotype quality of 30 were discarded (Figure 2A).

**Figure 2.**
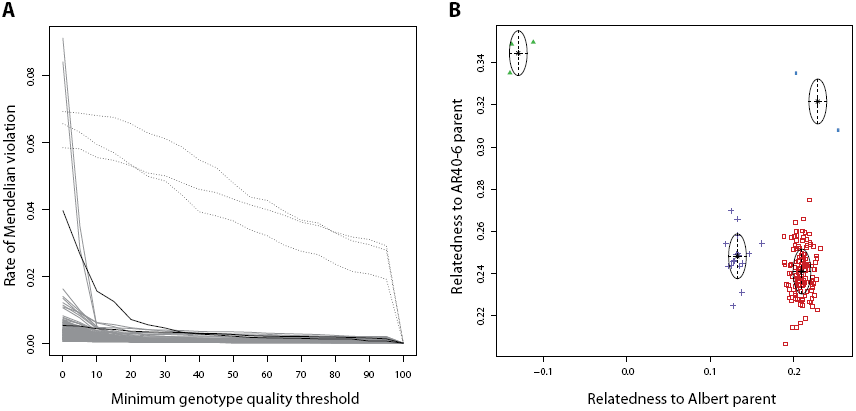
Analysis of parentage and sibling relationships. (A) The fraction of non-Mendelian genotypes detected in each individual in a population is plotted as a function of the minimum genotype quality (GQ) threshold. Off-types can be detected by a consistently high rate of Mendelian violation (black dotted lines), while, for legitimate progeny (solid gray), the non-Mendelian genotype rate is consistently lower than that of the putative parents (solid black lines). (B) A bivariate clustering analysis was performed on each population to verify parentage and full-sibling relationships. In this plot, we show the ARAL population as an example. Pairwise kinship coefficients *phi* are calculated between progeny and parents (MANICHAIKUL *et al.* 2010). Each putative progeny is represented as a point in two-dimensional space colored by its cluster assignment: full-sib F1 progeny are in red and S1 progeny of AR40-6 (therefore unrelated to Albert) are in green; individuals consistent with being progeny from a full sib of Albert crossed to AR40-6 are in purple; individuals consistent with being the progeny from an S1 of AR40-6 being crossed to Albert are in blue.

Secondly, a bivariate clustering analysis was performed with the Mclust (v4.2) R package using as the similarity measure the kinship coefficient *phi* calculated by vcftools (v0.1.12) (MANICHAIKUL *et al.* 2010; DANECEK *et al.* 2011; FRALEY *et al.* 2012; R CORE TEAM 2014). Values of *phi* were calculated between all individuals and the putative parents, each parent constituting an axis of potential genetic contribution (Figure 2B). Mclust calculates the optimal number of clusters under a Bayesian Information Criterion (BIC) regime for a given set of candidate multivariate Gaussian mixture models, the parameters for which are estimated via the Expectation-Maximization algorithm. Individuals are then classified into clusters by choosing the maximum conditional probability among the Gaussian distributions of the chosen mixture model (FRALEY *et al.* 2012). Individuals belonging to a single cluster, or several overlapping clusters, near (X,Y) = (0.25,0.25) were kept for downstream analysis. We performed a univariate analysis on the S1 population data because Mclust assumes that the two variables of the bivariate normal distribution do not have covariance near unity, and performing a bivariate analysis on an S1 population would violate this assumption.

### Calculating genetic maps for single populations

A genetic map was produced for each F1 mapping population using JoinMap software to estimate linkage, map order, and distance. To make mapping calculations more tractable, only one marker was retained from a set of markers at the same genetic position and individuals with over 50% missing data were removed. For each population, the minimum LOD threshold for grouping was determined by identifying grouping tree branches with stable marker numbers over increasing consecutive LOD values. Groups with three or more markers at the chosen minimum LOD threshold were retained for mapping. Specific LOD thresholds applied to each map are available in Table 4. Marker order and distances were determined using JoinMap’s Maximum Likelihood mapping algorithm for cross-pollinated (CP) populations. Default mapping parameters were assumed with the following modifications: spatial sampling thresholds reduced to 0.050, 0.025, 0.015, 0.010 and 0.005, and the number of Monte Carlo EM cycles increased to a value of 7. Maps were then compared to each other as a check for internal consistency (see next section) and genetically redundant markers that had been removed earlier were reincorporated into the component maps by assigning each marker to the LG and genetic position of the marker that was physically closest to it. The scaffolds in the draft assembly were used to determine physical distance. Finally, each map was converted from Haldane to Kosambi map units prior to merging.

**Table 4.** Mapping parameters and statistics. For each population and the composite map, we report the total number of markers and the number of markers used by JoinMap for map estimation, the minimum LOD threshold for defining a LG, and the number of LGs output from the grouping procedure. n.a. not applicable.

| Population | Number of markers | Number of framework markers | LOD threshold | LGs |
| --- | --- | --- | --- | --- |
| ARAL | 6,765 | 3,657 | 13.0 | 21 |
| KAR | 3,047 | 1,952 | 8.0 | 19 |
| MP4 | 3,392 | 1,903 | 6.0 | 19 |
| MP5 | 3,388 | 1,803 | 6.0 | 21 |
| MT | 3,991 | 2,301 | 10.0 | 18 |
| NCAR | 5,192 | 2,894 | 16.0 | 20 |
| NDLAR | 4,460 | 2,385 | 12.0 | 20 |
| NxA | 3,940 | 2,241 | 15.0 | 18 |
| TMEB419-S1 | 4,340 | 1,943 | 10.0 | 21 |
| 412x425 | 5,942 | 2,975 | 7.0 | 18 |
| **Composite map** | **22,403** | **n.a.** | **n.a.** | **18** |

### Quality control of component maps

Maps containing inter-marker genetic distances of 50 cM or greater were remapped with increased simulated annealing chain length and stopping chains. This process was iterated until maps no longer contained such distances or JoinMap ran out of memory. If these distances could not be removed before memory failure, individual markers near the interval with extreme values of “Nearest Neighbor Fit” (a measure reported by JoinMap to indicate the compatibility of one marker between two flanking markers) were removed by trial and error (recalculating the genetic map after each iteration) (removed markers are listed in File S1). If this failed to fix the gap, the LG was split manually into two sets of parent-specific markers (*i.e.* lm x ll and nn x np). Separate linkage maps were calculated for each parent using the methods described above. Single-parent maps that broke at the location of the initial marker gap were included for merging.

The relative correspondences and orientations of component LGs between populations were determined by examining dotplots, plotting the genetic positions of markers between pairs of maps (Figure 3). LGs were split or joined by following majority rule of the set of cognate LGs. For example, a component LG sharing markers with two LGs in each of the other component maps was split; multiple smaller LGs in one component map sharing markers with a single LG in each of the other component maps were joined. In joining, an interval was inserted equal to the average genetic distance found in the other corresponding component groups.

**Figure 3.**
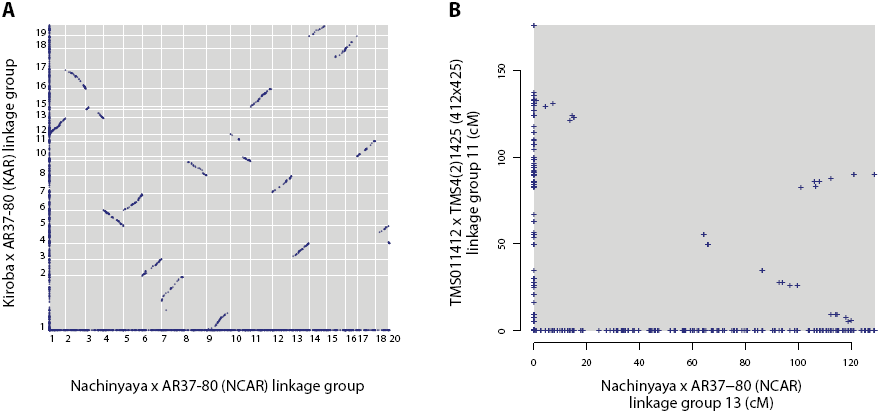
Pairwise comparison of single population maps. (A) An example of a pairwise comparison between maps. Every marker is plotted at a position corresponding to its genetic distance in each map (for shared markers), or along the axis (for markers unique to one map). Shared markers hence reveal the correspondences and relative orientations between LGs produced by JoinMap (arbitrarily numbered). Runs of shared markers appearing as roughly straight, continuous lines demonstrate marker orders consistent between the two maps; a positive or negative gradient indicates identical or opposite orientation, respectively, in the two maps. In addition, LGs to be joined or split are revealed, respectively, as multiple unusually small LGs in one map corresponding to a single LG in another map (NCAR LG 3, and KAR LGs 13 and 14) and as a single unusually large LG in one map corresponding to multiple LGs in the other (NCAR LGs 7 and 9, and KAR LG 1). (B) An example of a ‘V’-shaped dotplot pattern, typically observed in LGs that required (and could be corrected with) JoinMap parameters that increased the sensitivity (see Materials and Methods).

### Merging maps with LPmerge

We chose the ARAL map to be the reference for numbering and orienting LGs during merging (Table S1) because it was deeply sequenced and this map had the best agreement between genetic and physical marker order. The initial numbering of LGs followed that produced by JoinMap. As described above (Figure 3), a set of corresponding LGs and orientations was constructed for each of the 18 chromosomes in cassava. Each of these sets of corresponding LGs was merged into a composite map with LPmerge v. 1.4 (ENDELMAN AND PLOMION 2014), with maps being weighted by population size. LPmerge was run ten times with the maximum interval parameter set to each of the values in the range 1–10. We then chose the merged map with the value of maximum interval that gave the shortest total composite map length, as described in the documentation for LPmerge. For most LGs, the value of maximum interval was 1 (Table S1).

Each merged map was plotted along with all its component maps on a plot of marker number against genetic map distance. We noticed that markers at the end of some LGs (see *e.g.* Supplemental Figure 2) were tens of cM distant from adjacent markers. These markers were found to be present in only one contributing map and were distorting the merged map distances calculated by LPmerge. To fix this, terminal markers from single contributing maps were removed and another round of merging with LPmerge was performed. This process was repeated until there were no singleton markers at the end of a LG.

### Correcting scaffolding mis-assemblies

Mapped markers from all ten maps were considered simultaneously. Scaffolds from the v4.1 assembly (PROCHNIK *et al.* 2012) containing markers that mapped to different LGs were broken as long as the markers on the different LGs were derived from at least two maps or there were at least two markers from a single map. Mean pairwise linkage disequilibrium (LD) *r*^2^ statistics (PURCELL *et al.* 2007) were calculated using vcftools (DANECEK *et al.* 2011) for all variants derived from putatively mis-assembled scaffolds. Scaffolds were broken at scaffolding gaps within each region flanked by markers of different LG assignments if no evidence of LD was observed. If multiple scaffolding (*i.e.* sequence) gaps existed between flanking markers, the gap with the fewest supporting mate-pairs used for scaffolding the v4.1 assembly was broken. If multiple, equally supported gaps existed, the scaffold was broken at the largest gap. Minority markers from unbroken candidate mis-assembled scaffolds were removed.

### Scaffold anchoring and orienting

After breaking mis-assembled scaffolds, the v4.1 genome assembly was anchored to the genetic map and joined with 1000 Ns to produce pseudomolecules. Scaffold order was determined by median genetic position and orientation by the sign of Kendall’s rank correlation coefficient (*tau*) between physical and genetic positions. Scaffolds that could not be assigned to any LG (due to ambiguity in grouping or lack of markers) were not incorporated into the pseudomolecules and retained their original identifiers from the v4.1 assembly. Scaffolds that were anchored but could not be oriented were incorporated in their original (*i.e.* arbitrary) orientations. The LGs were then ordered by decreasing genetic size and numbered with Roman numerals to provide a canonical nomenclature (Figure 4).

**Figure 4.**
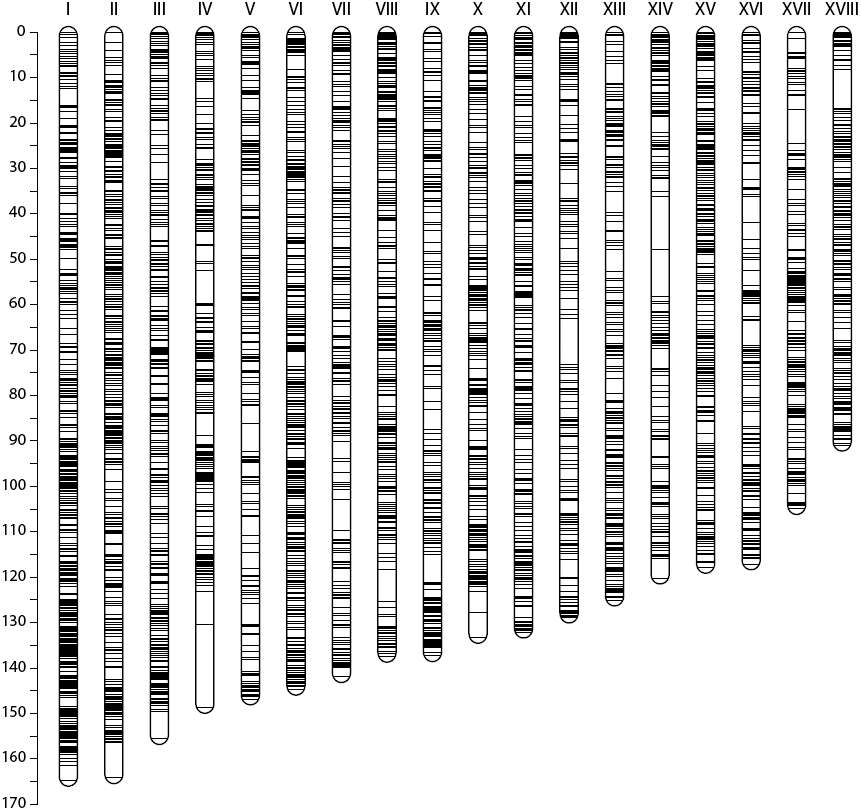
Marker distribution in the composite map. The scale on the left shows the map distance in cM. Our composite map consists of 18 LGs with a marker density of 1.95 non-redundant markers per cM. Previous maps did not recapitulate 18 LGs with a clear one-to-one mapping to our map, so we adopted a Roman numeral numbering system, ordered by decreasing genetic size.

## RESULTS AND DISCUSSION

To produce a robust and dense genetic map that can be used to follow traits segregating in diverse mapping populations, we generated ten mapping populations (Table 1) and genotyped them using a modified GBS approach (ELSHIRE *et al.* 2011). We were stringent in selecting markers, in order to both mitigate the effects of missing data inherent in a GBS approach and to ensure that variants were segregating in a Mendelian fashion. We then generated ten independent genetic maps from these populations. Comparison between them allowed us to identify cases where the standard map estimation approach was unable to resolve consistent LGs, and reconciling these differences produced a highly concordant set of maps. After constructing maps using a reduced set of stringently selected markers, we reintroduced genetically redundant markers into the maps, and these individual component maps were then integrated into a single composite framework map with the expected 18 LGs. Finally, we used this map to organize the v4.1 draft cassava genome sequence (PROCHNIK *et al.* 2012) into 18 pseudomolecules, anchoring 90.7% of the predicted genes onto LGs.

### Mapping populations

We analyzed nine F1 and one S1 mapping populations derived from 14 diverse cultivated varieties of cassava segregating for agriculturally important traits (Table 1). One population included here was recently used for a QTL analysis for cassava mosaic disease (CMD) resistance (RABBI *et al.* 2014b). Due to their desirable agricultural properties, some cultivars were used as parents in multiple crosses. Six populations from the Ferguson group at IITA were selected specifically for segregation of resistance to cassava brown streak disease. Several of these parents have a known wild species component in their pedigree (Table 2).

### Reduced representation sequencing

Several groups have developed inexpensive methods for generating a reduced representation of a genome with a restriction enzyme and sequencing to obtain genotypes (genotyping-by-sequencing or GBS) (ALTSHULER *et al.* 2000; BAIRD *et al.* 2008; DAVEY *et al.* 2011; ELSHIRE *et al.* 2011). To generate mapping markers in cassava, we adapted the GBS approach of Elshire *et al.* (2011) (Materials and Methods). We used Y-shaped GBS adapters to ensure that all pieces of DNA with adapter ligated to both ends could cluster and be sequenced. Since the same barcode was added to both ends of the DNA extracted from a given individual (see Figure S1), checking for the identity of barcodes from both ends provided additional quality control. To save time, we dispensed with the DNA and adapter drying step before digestion. To allow for robust ligation we phosphorylated the “reverse” oligos of our adapters at the 5’ end (Figure S1); although this can increase the number of adapter dimers, these dimers are effectively removed by size selection as noted below. To maximize the number of DNA fragments that are ligated to adapters, we used adapters in excess. This also prevents the formation of DNA concatemers, which would confound downstream analysis of reads. To increase the accuracy of mapping reads to the genome, the probability of detecting variation adjacent to a cut site, and the amount of data, we performed 100 bp paired-end sequencing, rather than the single-end 86 bp sequencing of Elshire *et al.* (2011).

During protocol optimization, we found that a gel-based size selection step amplified the desired size fraction more robustly. Fragments of sizes 400–800 bp cluster well on the flow cell yet are unlikely to contain adapter-mers. A gel-based size selection also provides for modularity in GBS approach: excising a narrower size range from the gel can be a simple way to sample fewer sites in the genome if fewer markers or more depth per site are needed, and/or to facilitate multiplexing more samples per lane. In the PCR, we used Phusion polymerase rather than *Taq* to minimize error, and we found that increasing the amount of PCR primer yielded more amplified library. We reduced the number of PCR cycles to five in order to reduce bias that can interfere with accurate variant calling. The 0.7x ratio of SPRI beads to DNA removes fragments less than 300 bp in length (QUAIL *et al.* 2008; LENNON *et al.* 2010), and thus effectively removes any remaining adapter-mers and excess PCR primer (neither of which are detected in appreciable amounts by Bioanalyzer on completed libraries).

### Genotyping-by-sequencing performance in cassava

Our adapted GBS protocol makes use of the *Ape*KI restriction enzyme used in the protocol developed by Elshire *et al.* (2011) and recommended by Hamblin & Rabbi (2014) for cassava. This enzyme and our method of library preparation allowed us to sample, per population, an estimated 85.5k restriction cut sites (with ten reads or more) distributed throughout the genome on 5,900 scaffolds (approximately 173 cut sites/Mbp) and sampling roughly 42K variable loci. Multiplexing up to 96 samples per lane of sequencing was effective for our outbred populations: we obtained an average of roughly 3.7M reads per individual in a typical 96-plex sequencing run.

### Genotype calling, filtering, and quality control

We genotyped all individuals against the currently available v4.1 draft cassava genome assembly with the GATK (MCKENNA *et al.* 2010). Based on an examination of the relationship between minimum genotype quality (GQ) threshold and the rate of violation of Mendelian segregation, we set a minimum GQ of 30, which effectively filtered low-confidence genotype calls (Figure 2A). In addition, we removed 44 individuals with excessive genotyping error or in which the pattern of marker segregation was inconsistent with that expected in an F1 or S1 population.

Since botanical seed-based fecundity is low in cassava, generating large mapping populations often requires crossing multiple cloned parental genotypes over an extended period of time, increasing the likelihood of error in tracking parental or progeny genotypes. We therefore performed a relatedness analysis (Materials and Methods) to identify and remove individuals that were not part of the intended cross (Figure 2B). Across the ten populations, we found 14 individuals that were unrelated to both parents, 68 related to only a single parent (26 selfs, 2 clones, 40 half-sib offspring), and identified 119 additional individuals with more complex relationships not suitable for mapping. Individuals identified by the relatedness and/or the Mendelian violation analyses were removed from further analysis. The number of progeny sequenced and used for map construction from each population is listed in Table 1. In one extreme case, none of the offspring in the MT population were related to the pollen parent that was sequenced (putatively TMS4(2)1425) (Figure S3) and furthermore neither these offspring nor the putative TMS4(2)1425 parent matched the TMS4(2)1425 parent of the 412x425 population. We could, however, confirm that most members of the MT population were full sibs. Given this fact, it was nevertheless possible to statistically infer reliable genotypes for the true pollen parent and estimate the combined genetic map for the population, as was performed for all other populations included in this study (see next section). Each population of verified full-sib progeny was then subjected to a significance threshold (P < 0.05) for deviation from Mendelian genotype frequencies, from which we obtained roughly 4,400 segregating sites per population (Table 4).

### Calculating genetic maps

Genetic maps were estimated for each population using the Maximum Likelihood mapping algorithm for CP populations implemented by JoinMap (v4.1). Forty-four (22.6%) component LGs contained intervals 50 cM or larger that could not be corrected by increasing the simulated annealing parameters and required manual curation. In this curation, markers that were causing the large gaps were identified by trial-and-error and removed. A total of 179 (out of 24,403 mapped) markers were discarded (File S1). After this step, one LG still contained a large gap. For this LG, a separate map was made from the markers unique to each parent and the LGs split into two pieces at the position of the gap. Subsequently, shared markers were reincorporated based on their scaffold coordinates.

### Resolving discrepancies between maps, refining maps, and gauging accuracy

Comparing the ten component linkage maps with each other allowed us to identify and correct inconsistencies including incorrectly split or joined LGs. We were able to verify long-range colinearity as the median number of markers shared between any two maps was 793 (quartiles Q1=656, Q3=983), or approximately 44 per LG (Figure 3A). Occasionally, the dotplot pattern of a LG was ‘V’-shaped (Figure 3B), indicating inconsistent marker order (*e.g.* a mapping error or a possible inversion) in one of the maps. In all but two cases (see below), the ‘V’ shapes could be resolved into linear relationships by increasing the simulated annealing parameters.

Most LGs had a one-to-one correspondence with an LG in each of the other populations. However, using a majority rule approach, it was clear that two component LGs required splitting and nine joins were needed. In one additional case, the decision to break or join was ambiguous, as five populations each contained two LGs that together corresponded to single LGs in the other five populations. We resolved this ambiguity by jointly examining the dotplots for the two-LG component maps against a corresponding single-LG map. From this, we discovered that none of the breaks in the different two-LG component maps occurred at the same genetic position. It is therefore likely that the LGs should be joined in the two-LG component maps to generate single LGs. Finally, three component LGs were not included in the merging because they could not be resolved into a map with a linear relationship with the corresponding LG from the other maps, and four were discarded because, although they could be matched with LGs in other maps, they contained too few markers to be confidently oriented.

Comparing maps to one another using dotplots provides a visual measure of internal consistency, while accuracy of marker order may be estimated by comparison to the physical sequence. We calculated Kendall’s rank correlation coefficient (*tau*) between physical and genetic positions on scaffolds that contained 10 or more markers and were not broken (described below). Across all maps and scaffolds examined, the median value *tau* was 0.84 (quartiles Q1 = 0.748, Q3 = 0.908), indicating good agreement between physical and genetic positions.

### Reincorporation of genetically redundant markers

In order to more fully represent the genetic diversity found within the mapping populations, 8,418 markers that had been excluded earlier based on their genetic redundancy (Materials and Methods) were reincorporated into the composite map. By creating datasets with the redundant markers included before and after map integration, and then comparing their median Kendall’s rank correlation coefficients *tau* (for 46 scaffolds with 50 or more markers, totaling 64.5 Mb of sequence), we found that including the redundant markers prior to map integration increased the marker order accuracy (*tau* = 0.765 vs. *tau* = 0.726). This improvement arises because the same genetically redundant marker is not always chosen to represent its genetic position in all component maps, and reincorporating redundant such markers increases the number of marker order constraints used by LPmerge (ENDELMAN AND PLOMION 2014) when a marker order conflict is encountered during map integration.

### An integrated framework map

With the exception of the three LGs noted above, the component maps were colinear with each other (*e.g.* Figure 3A). We next combined them into a single composite map. Genetic maps are merged by finding markers that are shared between the individual maps. Ideally all shared markers will be colinear, however in many situations discrepancies need to be resolved. Sometimes these discrepancies could be due to rearrangements in the genome of one of the parents, but most often are due to inaccurate genetic distances between markers because of insufficient recombinations, the stochastic nature of recombination, and errors introduced in individual map construction by missing and/or erroneous marker data. We used LPmerge (ENDELMAN AND PLOMION 2014) to generate a composite map from the ten component maps we had generated because it merges maps without recalculating recombination frequencies, drastically reducing the computational time required. LPmerge generates a consensus map with the minimum absolute error relative to the component maps, while preserving the marker order by imposing linear inequality constraints and deleting a minimum number of constraints if a marker order conflict is observed (ENDELMAN AND PLOMION 2014).

Combining maps from multiple independent crosses has the advantage of increasing the genetic diversity that is captured in the map; increasing support for marker order and position; and allowing markers from a single map to be placed relative to other markers. Composite maps have been generated from six Diversity Array Technology [DArT]-and RFLP-based maps for sorghum (MACE *et al.* 2009), six SSR-based maps for pigeonpea (BOHRA *et al.* 2012), and eleven SSR-based maps for groundnut (GAUTAMI *et al.* 2012) but, to our knowledge, no attempt has been made to integrate this many GBS-SNP based genetic maps.

After merging, we noticed that markers at the ends of LGs that belonged to a single map were placed on the merged map tens of cM from their neighbors (Figure S2). To improve the merging, we removed terminal markers belonging to a single map and repeated the merging until no further singleton markers remained. This removed a total of 182 markers (Table S1) to generate a 2,412 cM map containing 22,403 markers, averaging 134 cM and 1,245 markers per LG (Table 5, File S2). The SNP density (∼9 SNPs per cM) is substantially higher than that of the previous densest map (mean inter-SNP distance = 0.52 cM, (RABBI *et al.* 2014b), although this straightforward statistic does not account for the fact that many of the markers are genetically redundant (Table 5). For the first time, a cassava genetic map recapitulates the coherent set of 18 LGs matching the cassava chromosomes (Figure 4).

**Table 5.** Linkage groups in the composite map. The number of markers and genetic distances are shown for the 18 LGs. The average marker density is calculated for genetically non-redundant markers only *i.e.* only one marker at a given genetic position was included.

| Chromosome (LG) | Number of markers | Length (cM) | Average marker density (markers/cM) | Maximum inter-marker distance (cM) | Number of scaffolds anchored | Number of bases anchored |
| --- | --- | --- | --- | --- | --- | --- |
| I | 2,323 | 164.78 | 2.72 | 3.63 | 90 | 26,714,966 |
| II | 1,366 | 164.22 | 2.40 | 7.85 | 81 | 24,343,195 |
| III | 1,326 | 155.60 | 2.04 | 5.97 | 116 | 22,858,152 |
| IV | 1,459 | 148.73 | 1.64 | 18.41 | 93 | 21,649,806 |
| V | 1,330 | 146.87 | 1.57 | 6.12 | 81 | 23,125,959 |
| VI | 1,462 | 144.73 | 2.56 | 3.42 | 79 | 22,319,908 |
| VII | 848 | 141.96 | 1.65 | 6.73 | 88 | 18,888,952 |
| VIII | 1,212 | 137.48 | 2.01 | 7.21 | 111 | 23,111,533 |
| IX | 1,207 | 137.35 | 1.68 | 6.01 | 92 | 20,667,830 |
| X | 1,011 | 133.31 | 2.03 | 5.58 | 95 | 20,387,986 |
| XI | 1,330 | 132.16 | 2.12 | 2.56 | 96 | 20,727,479 |
| XII | 863 | 128.85 | 1.30 | 10.09 | 103 | 22,667,256 |
| XIII | 865 | 125.09 | 1.89 | 4.92 | 107 | 20,100,115 |
| XIV | 1,346 | 120.26 | 1.48 | 11.74 | 88 | 17,859,824 |
| XV | 1,548 | 117.89 | 2.54 | 2.56 | 72 | 20,107,995 |
| XVI | 920 | 117.13 | 1.56 | 5.71 | 75 | 18,834,636 |
| XVII | 974 | 104.95 | 1.71 | 7.48 | 95 | 20,464,108 |
| XVIII | 1,013 | 90.99 | 2.41 | 8.46 | 90 | 17,820,998 |
| **Total** | **22,403** | **2,412.35** | **35.31** | **124.45** | **1,652** | **382,650,698** |
| **Average** | **1,245** | **134.02** | **1.96** | **6.91** | **91.78** | **21,258,372** |

We observed a steady increase (albeit with diminishing returns) in the number of markers and amount of assembled sequence that can be placed as a function of the number of mapping populations that were used (Figure 5). We note, however, that even with the addition of the tenth component map, we were still adding more markers and able to anchor more genome sequence (Figure 5). Future work will merge additional maps to take advantage of this feature. However, because the majority of markers were present in only one map (12,474 out of 22,403) they may be placed inaccurately in the merged map as they are not constrained by more than one map. This will be improved upon the generation of a less-fragmented genome assembly.

**Figure 5.**
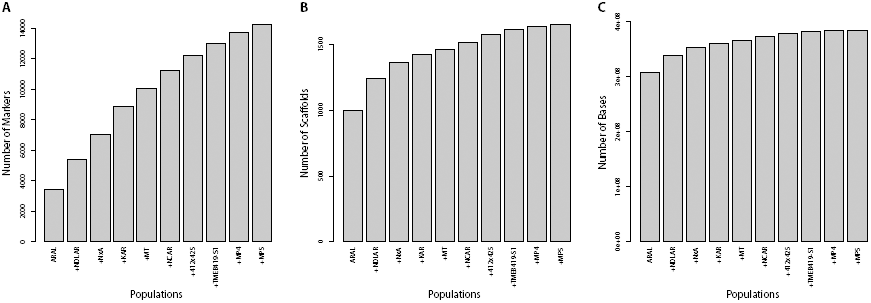
Additional maps incorporate more markers, scaffolds, and anchored bases. These plots show the effects of adding maps to the framework map. Each additional map incorporates more genetically non-redundant markers (A) into the framework map, but the number of scaffolds incorporated is saturating (B) and the number of mapped bases (C) is reaching a plateau. This is because the scaffolds being added in later maps are getting smaller and smaller and hence adding ever fewer bases.

All maps were to a large extent colinear with the integrated framework map (Figure S4), however, component LGs from NxA (see *e.g.* Chromosome II, Figure S4B) as well as 412×425 (see *e.g.* Chromosome XVII, Figure S4Q) displayed notable divergence from this, likely because these maps were of lower quality. Lastly, across the whole genome, we observed four gaps larger than 10 cM (terminal portion of Chromosome V; two on Chromosome III and one on Chromosome VIII). These could be recombination hotspots or regions that are identical-by-descent and thus lack polymorphisms.

### Generation of chromosome pseudomolecules

One of our primary goals was to organize the fragmented draft genome assembly (PROCHNIK *et al.* 2012) into chromosome-scale sequence, therefore it was necessary to first detect possible discrepancies between the genetic map and physical sequence. Of the 1,347 scaffolds that contained multiple markers (median size = 174 kb), 41 scaffolds contained markers along their lengths that mapped to different LGs. Since all component maps used to generate the composite map were internally consistent, these discrepancies were unlikely to be due to errors in the genetic map and suggested a sequence mis-assembly in the scaffolds in question. Manual review, along with the aid of linkage disequilibrium and scaffolding information, allowed us to break likely misjoins where weak scaffolding linkages had been made, often in regions with a high density of scaffolding gaps. After breaking, we then organized the sequence assembly into pseudomolecules by anchoring the scaffolds onto LGs using their median genetic position. The resulting chromosome-scale sequences incorporate 1,608 scaffolds and 382 Mb (71.9%) of the v4.1 draft genome sequence (Table 5). This includes 462 (94.9%) of the N50 scaffolds, 1,430 (53.9%) of the N90 scaffolds, and 27,825 (90.7%) of the 30,666 predicted protein-coding genes. Of the scaffolds anchored onto the assembly, we could also orient 1,024 (63.7%) of them by calculating the Kendall rank correlation coefficient *tau* between physical and genetic positions. Together, these oriented scaffolds comprise 315 Mb of sequence.

### Concluding remarks

Here we present a consensus genetic map of cassava that combines 10 mapping populations. Unlike many previous maps (FREGENE *et al.* 1997; OKOGBENIN *et al.* 2006; KUNKEAW *et al.* 2010; SRAPHET *et al.* 2011; RABBI *et al.* 2012; RABBI *et al.* 2014a; RABBI *et al.* 2014b) we recovered the 18 LGs predicted by the karyotype of *M. esculenta* (DE CARVALHO AND GUERRA 2002) and at the same time dramatically increased the number of markers to over 22,400. The vast majority of genes in the current draft genome were placed on a LG and at their approximate chromosomal position, resulting in the first chromosome-scale assembly of cassava and a map that will serve as a valuable guide for future genome assembly improvements. Our approach can be readily extended to include future mapping populations, although as seen here care must be taken to explore discrepancies between biparental or selfed maps that may arise from mistaken parentage, missing or erroneous data, and the details of map construction. Using multiple maps at the same time allows such individual component map problems to be identified and corrected (or excluded from merger).

Within the context of disease threat, climate change and food insecurity, improving cassava for smallholder farmers in developing countries is becoming more urgent. A number of projects and collaborations are already engaged in this task. These initiatives are rapidly incorporating modern genetic tools and techniques such as genomic selection and QTL analysis, techniques that depend on an accurate genetic map and genome assembly. The chromosome-scale genome sequence and composite map we report here will allow cassava geneticists and breeders to generate the robust cassava varieties needed both for food security and to improve the economic conditions of smallholder farmers around the world.

## ADDITIONAL INFORMATION

The composite genetic map is available in File S2, and also at CassavaBase (www.cassavabase.org). Chromosomal pseudomolecules are available at Phytozome (www.phytozome.org). All raw sequence reads will be available in the Short Read Archive.

## ACKNOWLEDGEMENTS

Richard Harland and Lisa Brunet, for early input into optimization of the GBS protocol. Craig Miller lab/Andrew Glazer for reagents and helpful discussion. Minyong Chung, Justin Choi, Karen Lundy, and other members of the VCGSL at UC Berkeley, for advice and technical assistance with sequencing. Jeff Endelman for advice on using LPmerge. Ed Buckler, Jean-Luc Jannink, and Martha Hamblin, for helpful discussion.

JBL, JVB, CMH, and work at UC Berkeley were funded by Bill and Melinda Gates Foundation (BMGF) Grant OPPGD1493 to SR, DSR, and the University of Arizona. Development of populations at IITA Ibadan was funded by the CGIAR Research Program on Roots, Tubers and Bananas. Next Generation Cassava Breeding grant OPP1048542 from BMGF and the United Kingdom Department for International Development supported SEP and work at NRCRI and IITA. Development of the mapping populations in Tanzania was funded by BMGF grant OPPGD1016 to IITA. The work conducted by the U.S. Department of Energy Joint Genome Institute is supported by the Office of Science of the U.S. Department of Energy under Contract No. DE-AC02-05CH11231. This work used the VCGSL at UC Berkeley, supported by National Institutes of Health S10 Instrumentation Grants S10RR029668 and S10RR027303.

